# An approach for coherent periodogram averaging of tilt-series data for improved CTF estimation

**DOI:** 10.1101/2024.10.10.617684

**Authors:** Sagar Khavnekar, William Wan

## Abstract

Cryo-electron microscopy (cryo-EM) has become an indispensable technique for determining three-dimensional structures of biological macromolecules. A critical aspect of achieving high-resolution cryo-EM reconstructions is accurately determining and correcting for the microscope’s contrast transfer function (CTF). The CTF introduces defocus-dependent distortions during imaging; if not properly accounted for, the CTF can distort features in and limit the resolution of 3D reconstructions. For tilt-series data used in cryo-electron tomography (cryo-ET), CTF estimation becomes even more challenging due to the tilt of the specimen, which introduces a defocus gradient across the field of view, as well as the low dose and signal in individual tilt images. Here, we describe a simple algorithm to improve the accuracy of CTF estimation of tilted images by leveraging the tilt-series alignment parameters determined for tomographic reconstruction to explicitly account for the tilted specimen geometry. In brief, each tilt image is divided into patches, each of which are then stretched according to their defocus shift. These are then summed to provide a coherent power spectra at the tilt axis, which can then be used in standard CTF estimation algorithms. This uses all the data in each image to enhance the visibility of Thon rings, thereby improving high-resolution CTF estimation and subsequent enhancements in the resolution of subtomogram averages.

## Introduction

Biomolecular structure determination using cryo-electron microscopy (cryo-EM) methods such as single particle analysis (SPA) or subtomogram averaging (STA) relies on averaging multiple copies of a repeating structure. Averaging is necessary to achieve adequate orientational sampling for 3D reconstruction, increase the signal to noise ratio (SNR) of high-resolution features, and overcome image aberrations from electron optics (Young & Villa, 2023; Turoňová & Wan, 2024; Watson & Bartesaghi, 2024). The primary aberration in cryo-EM images is the contrast transfer function (CTF) (Erickson & Klug, 1971), which is inherent to the spherical objective lenses used in electron microscopes. The CTF is strongly dependent on defocus, which is routinely applied to improve contrast in cryo-EM images (Erickson & Klug, 1971; Downing & Glaeser, 2008). The CTF causes sinusoidal amplitude modulations and phase oscillations in Fourier space that periodically inverts the contrast in certain frequency ranges, resulting in destructive interference and loss of resolution. This can be partially corrected for by applying a filter to “flip” the phases of inverted regions, allowing for constructive interference of higher resolution information, though the amplitude modulations of the CTF cannot be directly corrected (Downing & Glaeser, 2008). These amplitude modulations are instead normalized by averaging images with different defocus values. Imprecise CTF-correction can partially restore signals, but residual destructive interference is still present at higher resolutions (Zanetti *et al*., 2009). As such, 3D reconstructions from data that has not been accurately CTF-corrected can have residual real-space distortions and limited resolution. Therefore, accurate CTF estimation is imperative for high-resolution structure determination, which has led to the development of a number of packages to estimate CTF for cryo-EM micrographs (Mindell & Grigorieff, 2003; Rohou & Grigorieff, 2015; Zhang, 2016; Su, 2019; Elferich *et al*., 2024; Zhang *et al*., 2024).

From an image processing perspective, the CTF can be thought of as an image filter and is generally modelled as a 2-dimensional function of the spatial frequency vector **k**:

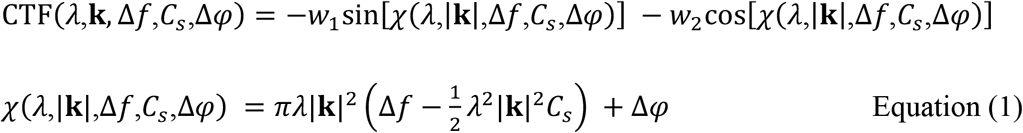

where the frequency-dependent phase shift *χ* is a function of the electron wavelength, Δ*f* is the objective defocus value, *C*_*S*_ is the spherical aberration of the objective lens, Δ*φ* is the additional phase shift introduced by a phase plate, *w*_2_ is the fraction of total contrast attributed to amplitude contrast, and *w*_1_ is the relative phase contrast given as 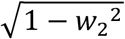 (Rohou & Grigorieff, 2015).

The CTF is typically estimated from the power spectra of a micrograph, where the amplitude modulations appear as set of concentric rings, referred to as Thon rings (Thon, 1966). Calculation of power spectra, either by periodogram averaging or Fourier-rescaling (Fernández *et al*., 1997; Rohou & Grigorieff, 2015), enhances the appearance of Thon rings compared to the raw Fourier transforms by minimizing Fourier-space noise. Most programs estimate the CTF by fitting Equation 1 to a background subtracted power spectra of the micrograph; subsequent CTF-correction is then performed by using the fitted CTF to determine the spatial frequences with a negative phase and making them positive in the image, either by multiplying by -1 (phase flipping), multiplying by the fitted CTF, or applying a Wiener filter (Downing & Glaeser, 2008).

The main variable that is fitted during CTF estimation is the defocus. Simply put, defocus is when the specimen does not sit in the focal plane of the objective lens. Practically, a combination of specimen thickness, nonplanar specimen geometry, or specimen tilt with respect to the optical axis results in different areas of the specimen can have different defocus values. When using power spectra calculated from the entire micrograph (global CTF estimation), higher resolution Thon ring signals can diminish due to the averaging of varying local CTFs across specimen. One of the earliest algorithms for estimating the CTF of tilted images is the CTFtilt algorithm (Mindell & Grigorieff, 2003) implemented in CTFFIND3 (Mindell & Grigorieff, 2003) and CTFFIND5 (Elferich *et al*., 2024). The initial implementation of CTFtilt estimates CTF values for tiles extracted along parallel lines while simultaneously varying the angle of these lines to determine the tilt axis angle. A correct tilt axis angle will have highest resolution Thon ring fitting across each parallel line, while the defocus values of each line can be used to determine the tilt angle. The recent version tiles an image in a square grid and fits the CTF while solving a geometric model that best fits all tiles. Warp estimates the tilt of images by tiling images and estimating the CTF from each tile while also applying a regularization parameter to estimate the geometry of the specimen. Both these approaches rely on fitting Thon rings to subsets of the full image, which generally requires a relatively high SNR for precise fitting. Tilt-series data have an additional challenge over tilted single particle data due to the fact that a tilt-series image often has an order of magnitude less electron dose, resulting in an extremely low SNR of both the image and its power spectra. As such, patch-based CTF estimation methods that only use portions of the image for local power spectra calculation further degrading the SNR, limiting the ability to accurately fit Thon rings. While patch based methods are necessary to simultaneously determine defocus and tilt geometry, this is arguably unnecessary in tilt-series data, as the tilt axis angle and the tilt angles for individual tilt images are determined during tilt series alignment (Mastronarde, 2006); this ‘*a priori*’ knowledge can then be directly used during CTF estimation.

Here we describe a simple algorithm for improving CTF estimation of tilted images by using tilt-series alignment information to generate a coherent periodogram average from a whole tilted image. We do this by first tiling each image and geometrically calculating the relative height of each tile from the tilt axis, providing a defocus offset value. By applying a defocus-dependent stretching factor to these patches, we generate a single power spectrum that represents the CTF on the specimen tilt axis. After calculation of the tilt-corrected periodogram average, the CTF can then be fitted using standard packages such as CTFFIND4 (Rohou & Grigorieff, 2015). This periodogram averaging approach is similar to those implemented in Bsoft (Heymann, 2021), Ctfplotter (Mastronarde, 2024), and CTFMeasure (Zhang *et al*., 2024). After averaging, Bsoft also fits CTF to 2D power spectra similar to CTFFIND, while Ctfplotter integrates a series of wedges into 1D power spectra to estimate defocus and astigmatism. CTFMeasure is an approach similar to Bsoft but includes additional algorithms for refining the tilt geometry.

We demonstrate that our periodogram averaging approach improves the appearance of higher-resolution Thon rings in power spectra from tilt-series data on both amorphous carbon films and cryo-focus ion beam (FIB) milled lamellae of *S*. cerevisiae; these improved Thon rings result in more accurate estimation of the CTF. To show the suitability of this approach for cellular cryo-ET and STA, we demonstrate that this improvement in CTF estimation also results in improved 80S ribosome subtomogram averaging using the EMPIAR-11756 dataset (Rangan *et al*., 2024; Wan *et al*., 2024).

### Tilt-corrected periodogram averaging

During tilt-series data collection (Figure 1A), the specimen is rotated around the microscope stage tilt-axis (red) at discrete angular steps (tilt angles, green). During tilt-series alignment, it is convention to rotate and shift the micrograph to place the microscope stage tilt-axis is on the central Y-axis of the image; as such, stage tilt is also referred to as ‘Y-tilt’. Additionally, the specimen is often not perfectly parallel with the plane of the microscope stage, resulting in an additional oblique tilt when the stage is at 0° tilt. This effectively results in additional tilt around the tilt-axis (Y-axis pretilt) as well as an additional tilt perpendicular to the tilt axis, (X-tilt, dark blue) (Figure 1A, right panel). Altogether, this results in differences in height across the specimen at each tilt and thereby a different defocus offset Δ*z*{*effective*} at every point **P** with respect to that at the origin **O** on the stage axis (Figure 1B). The projections of these points on the image plane are depicted as **P’** and **O’**, respectively. The goal of CTF estimation is to determine accurate CTF parameters at the origin. However, due to the gradient across the specimen plane when the specimen is tilted, summed power spectra Σ_*X*′_ *CTF*(*X*′) (**O’**,**P’**,**…** ⊂ **X’**) often results in diminishing thon rings. (Figure 2A,D).

**Figure 1.**
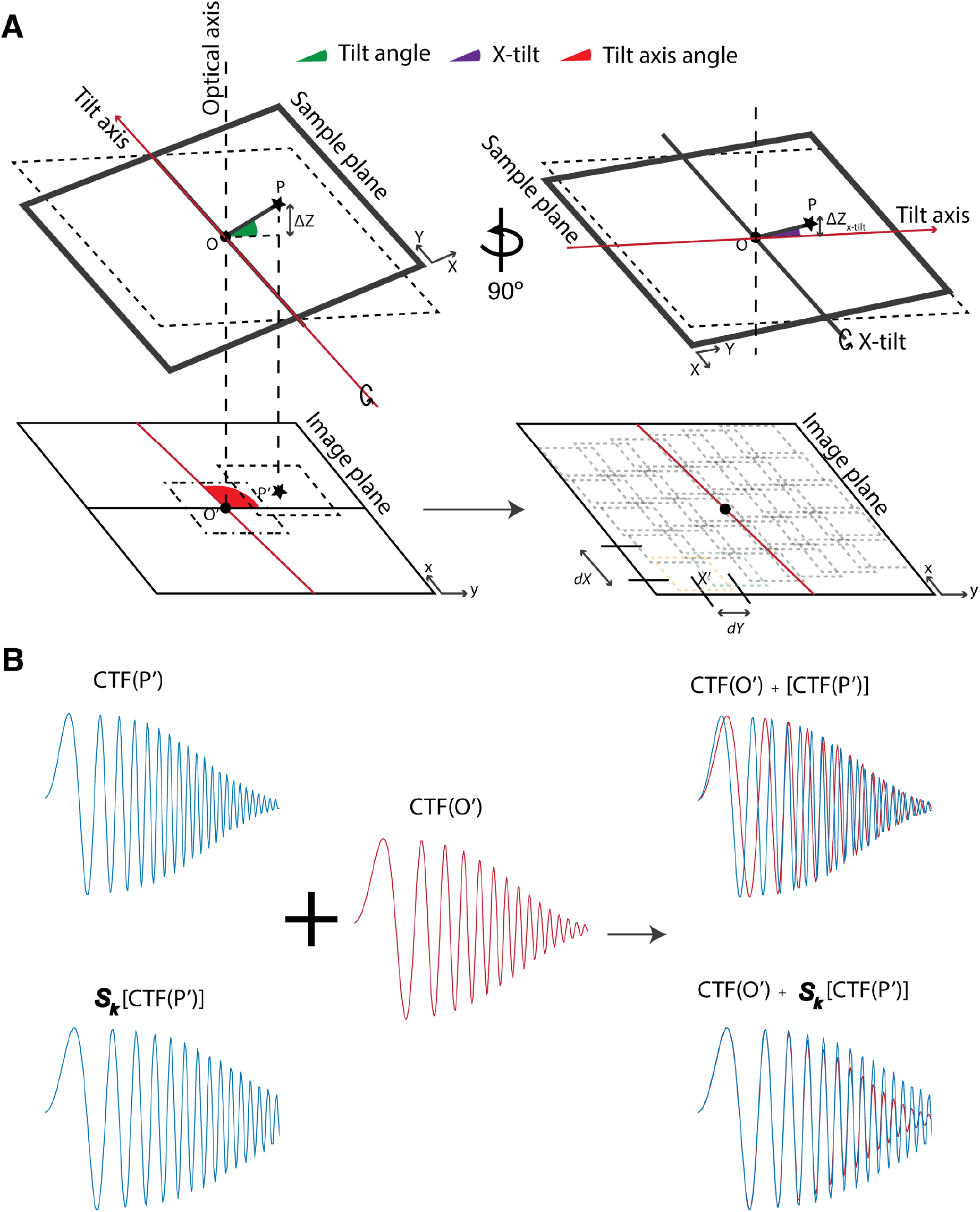
The geometry of tilted specimens and its effects on local CTF. (A) Relationship between specimen coordinate system and the image coordinate system. During tilt-series collection, the specimen is rotated along the tilt-axis (red) at discrete angular steps (tilt angles, green). The specimen often is often at an oblique angle as opposed to lying perfectly flat in the plane of the microscope stage.; this misorientation results in additional tilt with respect to the X axis in the plane of the microscope stage (x-tilt, dark blue). Altogether this results in a difference in height and hence defocus (Δ*z*) at point **P** with respect to that at the origin (**O**). The projection of these points on the image plane is depicted as **P’** and **O’**, respectively. In addition, the orientation of the tilt axis with respect to the Y axis of the image plane is given as the tilt-axis angle (red cone). For coherent periodogram averaging, tiles are extracted in a rectangular grid aligned with the tilt axis with a user-defined tile size. In the direction parallel to the tilt axis (dX) tiles are extracted with an offset of half the tile size, while in the direction perpendicular to the tilt axis (dY), tiles are extracted with a user-defined defocus offset. (B) The goal of CTF estimation of tilt-series data is to determine accurate CTF parameters at the origin. However, due to the defocus gradient across the specimen when it is tilted, summed power spectra Σ_*X*_′ *CTF*(*X*′) often results in diminishing Thon rings. **O’**,**P’**,**…** ⊂ **X’**. TiltCTF takes defocus gradient into account and applies a linear stretching operator ***S***_***k***_. The resulting summed power spectra, Σ_*X*_′ ***S***_***k***_[*CTF*(*X*′)] allows for accurate estimation of CTF parameters at the origin **O/O’**

**Figure 2.**
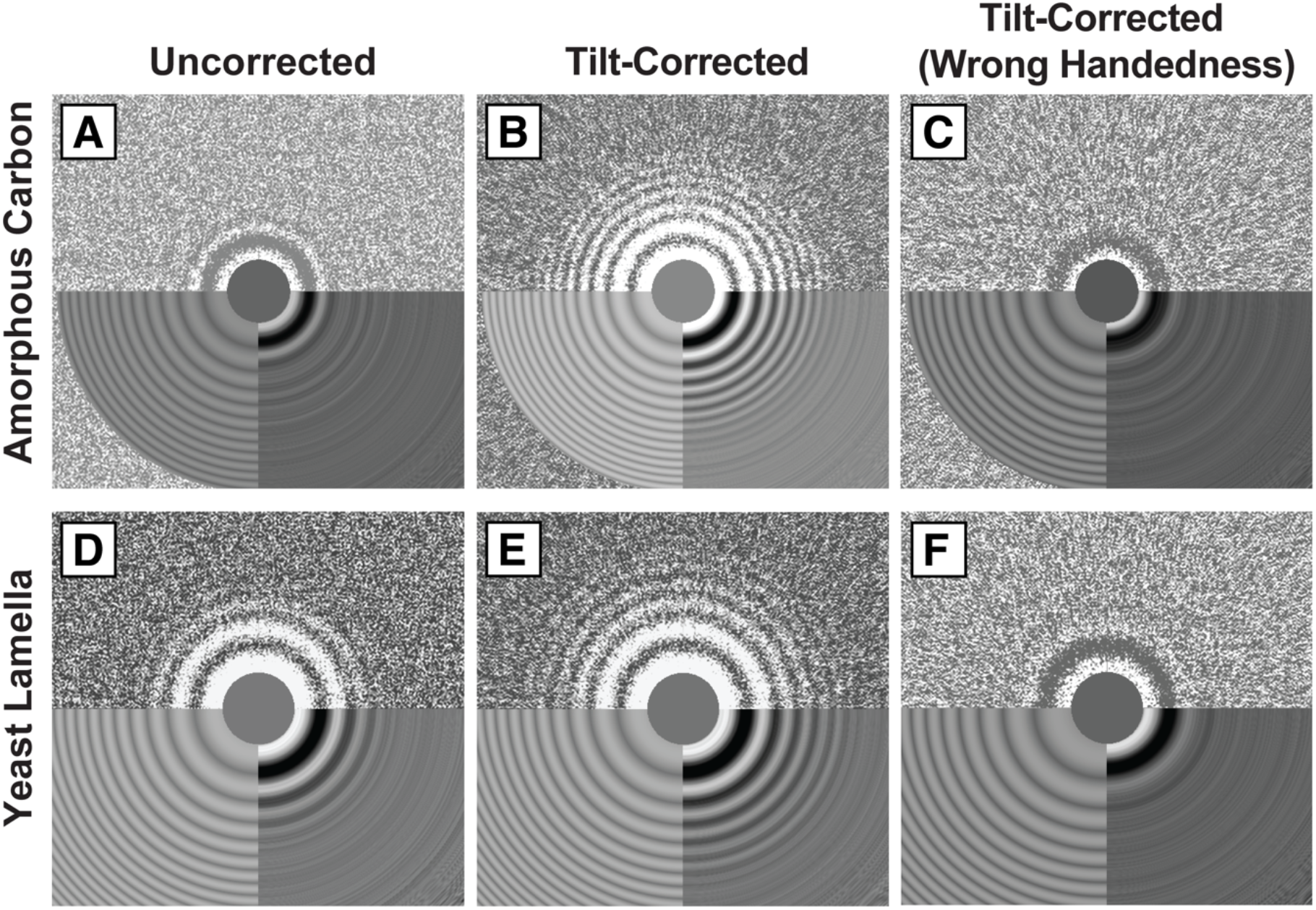
Tilt-corrected periodogram averaging improves Thon ring appearance and fitting at high tilts. CTF fitting is shown as summed power spectra (top half), fitted power spectra (lower left quadrant), and the fitted equiphase average (lower right quadrant). Fitting of an amorphous carbon specimen at 56º tilt angle using uncorrected (A), tilt-corrected (B), and tilt-corrected periodogram averaging with wrong tilt handedness (C). Without correction, Thon ring signal is lost by the second ring. Tilt-corrected periodogram averagings show fitting beyond the 9^th^ Thon ring, while correction with the wrong handed tilt results in loss of Thon rings. Fitting of *S. cerevisiae* cryo-FIB milled lamella specimen at -60 º tilt angle using uncorrected (D), tilt-corrected (E), and tilt-corrected periodogram averaging with wrong tilt handedness (F) for. Even for low-dose and thick specimens (∼200 nm), tilt-corrected periodogram averaging outperforms uncorrected periodogram averaging; in this example, fitting is up to the 7^th^ Thon ring.

Tilt-series alignment is the process of solving for the shifts and rotations in each tilt image in order to line up their common tilt axis prior to tomographic reconstruction. Tilt-series alignment methods typically track fiducial markers such as gold nanoparticles or specimen features within the tilt images, both of which rely on lower-resolution features that are largely independent of image defocus (Mastronarde & Held, 2017; Zheng *et al*., 2022). Within the field of view of a tilted specimen, the changes in the CTF profile due to defocus variations can largely be modelled as a linear stretching (Heymann, 2021; Zhang *et al*., 2024; Elferich *et al*., 2024; Mastronarde, 2024). As such, tilt-series alignment parameters can be used as *a priori* knowledge to calculate the difference in defocus between two points in the image, which is used to calculate the linear scaling factor ***S***_***k***_ required to match the two CTF profiles. By applying this scaling factor to Fourier transforms of patches with different defocus values, their CTF modulations can be summed constructively. In our approach, we calculate a periodogram average by tiling the specimen, calculating the defocus at the center of each tile using the tilt series alignment parameters, using that defocus value to calculate and apply a stretch factor ***S***_***k***_ to the Fourier transform of each tile, and summing all tiles. The resulting summed power spectra contains signal from all parts of the image (Figure 2B), Σ_*X*_′ ***S***_***k***_[*CTF*(*X*′)] allowing for the accurate estimation of CTF parameters at the origin **O/O’**.

### Tilt-corrected periodogram averaging improves thon ring fitting and subtomogram averaging

To demonstrate the ability of our algorithm to constructively average Thon rings in tilted specimens, we acquired high-dose (∼ 10 e^-^/ Å^2^) tilt-series on amorphous carbon. Without accounting for tilt (uncorrected), it is evident that the high-resolution Thon rings diminish at high tilt angles due to destructive interference (Figure 2A). When tilt-corrected periodogram averaging is performed, Thon rings are apparent to high resolution due to constructive interreference from each tile, allowing for the accurate estimation of CTF parameters (Figure 2 B). Alternatively, an incorrect handedness of the tilt angle results in even more destructive interference (Figure 2C). Similarly, tilt-corrected periodogram averaging improves the appearance of high-resolution Thon ring for low-dose tilt-series acquired on cryo-FIB milled lamella from vitrified S. cerevisiae (Figure 2D–F).

Accurate CTF estimation is of paramount importance for STA, as slight inaccuracies can result in significant loss of high-resolution information. To test the suitability of our tilt-corrected periodogram averaging algorithm for cellular cryo-ET and STA, we benchmarked out algorithm on the EMPIAR-11658 *S. cerevisiae* FIB-milled lamella dataset (Rangan *et al*., 2024; Wan *et al*., 2024). 3D CTF-corrected tomograms that used CTF estimates from global periodogram average resulted in a 14.3 Å subtomogram average of the 80S ribosome (Figure 3A,B). Using the same subtomogram alignment parameters, we then used our tilt-corrected periodogram averaging for CTF estimation followed by 3D CTF-correction, which resulted in 8.9 Å subtomogram average (Figure 3A,C). This improvement in resolution allowed us to perform further STA using an expanded low pass filter, resulting in a 8.4 Å subtomogram average (Figure 3A,D).

**Figure 3.**
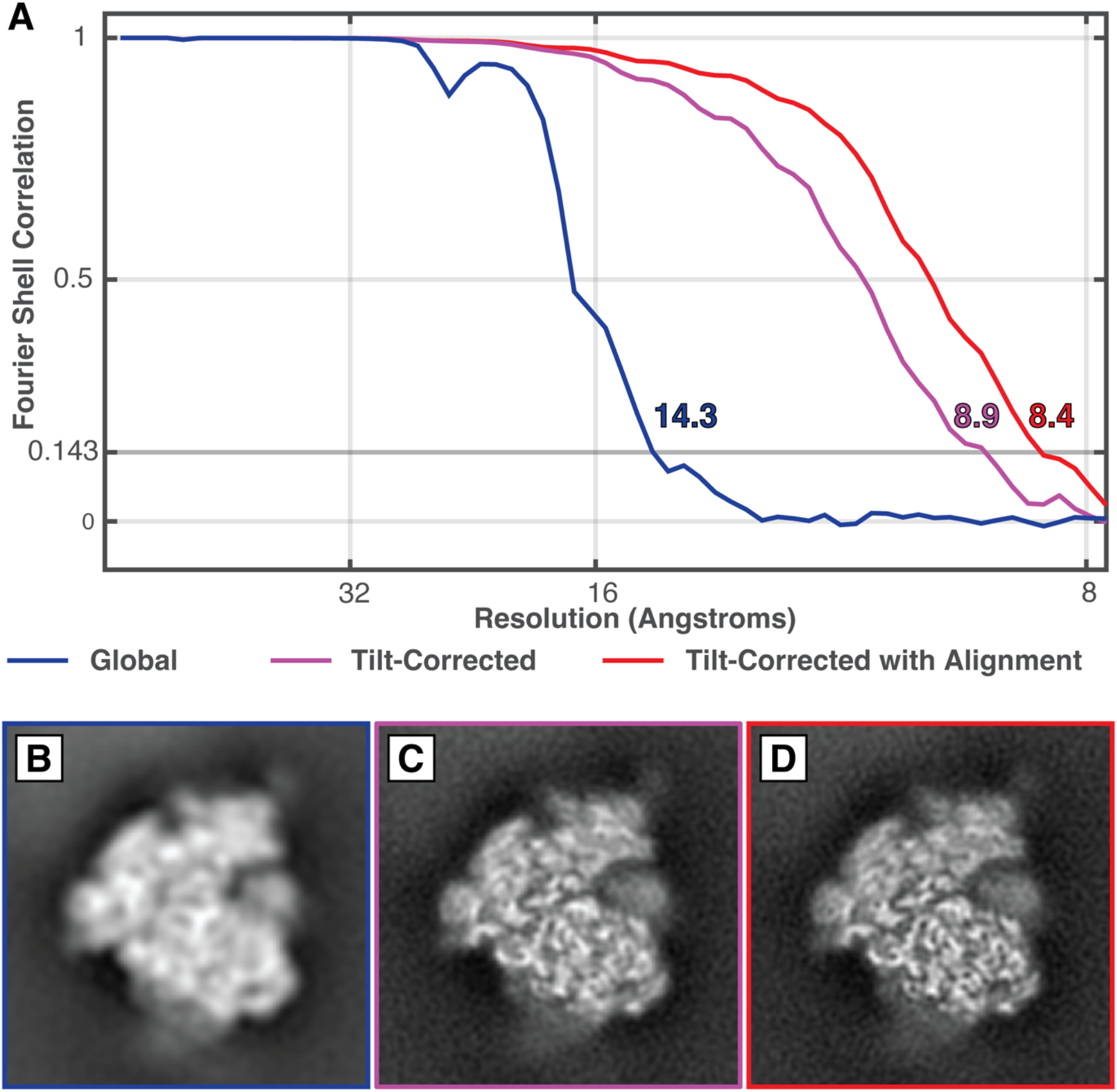
Accurate CTF estimation results in improved STA. A) “Gold-standard” Fourier shell correlation (FSC) plots for 80S ribosome subtomogram averages from EMPIAR-11658 dataset using STOPGAP and novaCTF 3D-CTF-corrected tomograms. CTF estimation was performed using CTFFIND4 uncorrected (blue) and tilt-corrected periodogram averages as implemented in tiltCTF (magenta). After averaging of tilt-corrected data, additional STA with a wider low pass filter further improved resolution (red). B–D). Orthographic slices through the STA maps plotted in A).

## Conclusion

Accurate estimation of CTF parameters is essential for restoring high-resolution features when using averaging methods such as SPA and STA. CTF estimation is performed by calculating the power spectrum of each image and fitting a CTF curve to the Thon rings in the spectra. For tilted images, this can be challenging as calculating a global power spectrum from the whole image results in a defocus-dependent dampening of high-resolution features due to destructive interference of varying CTF modulations. While this can be ameliorated by fitting power spectra calculated from local regions, this can be of limited effectiveness in tilt-series data. This is because the total dose in tilt images is typically an order of magnitude lower than for tilted SPA images resulting in reduced SNR, which is exacerbated when calculating local power spectra.

Here we present an algorithm for calculating tilt-corrected periodogram averages, which use the information from an entire image to calculate a power spectra that represents the CTF at the tilt axis. Our approach makes use of predetermined tilt-series alignment parameters to define the geometry of the specimen and apply appropriate stretch factors to the Fourier transforms of local tiles to calculate a constructive periodogram average from the entire image. Using tilt-series of amorphous carbon as well as FIB-milled cellular lamella, we show that our algorithm restores high-resolution Thon rings with sufficient intensity for accurate CTF fitting. We also show that the improved accuracy of CTF fitting improves subtomogram averages of 70s ribosomes from *S. cerevisiae*.

We have implemented this algorithm as the “tiltCTF” module in our TOMOMAN (Khavnekar *et al*., 2024) cryo-ET preprocessing package. TOMOMAN is an extensible software package that acts as a wrapper for various cryo-ET preprocessing tasks to allows users to customize their own processing pipelines with various internal algorithms and external software packages. The integration of tiltCTF into TOMOMAN allows users to transparently use the tilt-series alignment metadata from packages such as IMOD (Mastronarde & Held, 2017) or AreTomo (Zheng *et al*., 2022) to perform CTF estimation on tilt-series data. TiltCTF parses tilt-series alignment parameters stored in TOMOMAN’s internal metadata format to calculate tilt-corrected periodogram averages; these are then fitted using CTFFIND4. In keeping with TOMOMAN’s experimental ethos, decoupling of the periodogram averaging step from the fitting step potentially allows users to test different fitting algorithms, including those developed specifically for SPA. TiltCTF is written in MATLAB and provided as open-source code; TOMOMAN is available at https://github.com/wan-lab-vanderbilt/TOMOMAN.

## Acknowledgements

This work was supported by U.S. National Institutes of Health grant DP2GM146321 (to WW). WW is also a Pew Scholar in the Biomedical Sciences and supported by the Pew Charitable Trusts. This work was conducted in part using the resources of the Advanced Computing Center for Research and Education at Vanderbilt University, Nashville, TN. Some of this work was performed at the Max Planck Institute of Biochemistry with support from Wolfgang Baumeister, Juergen Plitzko, and John Briggs for resources and infrastructure. We would also like to thank Max Planck Computing and Data Facility (MPCDF) for the computational infrastructure. SK would like to thank the Max Planck Society and the International Max Planck Research School for graduate school funding.

## Methods

### Sample prep, acquisition, image processing

#### A. Carbon tilt-series

For the carbon tilt-series dataset, a 1:4 suspension of 3x concentrated 10 nm gold fiducials (Aurion) in water was applied onto a glow-discharged 200 mesh Quantifoil MultiA copper grid. Samples were vitrified in a liquid ethane/propane mixture using a Vitrobot Mark IV (Thermo Fisher Scientific).

Carbon tilt-series datasets were collected using a Thermo Scientific Titan Krios G3i equipped with a Selectris X energy filter and Falcon4 direct detector. Tilt-series were collected with a dose-symmetric tilt scheme (Hagen *et al*., 2017) using SerialEM (Mastronarde, 2005). Tilt range was ± 60º with 3º angular increments. Target focus was set to -3 μm. Tilt images were acquired in EER (Electron Event Registration) mode with a calibrated physical pixel size of 1.22 Å and total dose of 10 e^-^/Å^2^.

The data was preprocessed using TOMOMAN (Khavnekar *et al*., 2024). Motion correction was performed using RELION’s implementation of MOTIONCOR with EER support (Zivanov *et al*., 2019). Periodogram averages for uncorrected, tilt-corrected, and tilt-corrected with wrong handedness were calculated using tiltCTF module in TOMOMAN, which uses CTFFIND4 (Rohou & Grigorieff, 2015) for CTF fitting.

#### B. EMPIAR-11658

To demonstrate the improvements in subtomogram averaging from CTF estimation by tilt-corrected periodogram averaging, we processed *S. cerevisiae* ribosomes from EMPIAR-11658 using STOPGAP (Wan *et al*., 2024). 3D-CTF corrected tomograms were reconstructed using novaCTF (Turoňová *et al*., 2017) module of TOMOMAN, using CTF parameters from the minimal TOMOMAN project deposited as part of EMPIAR-11658. These CTF parameter were estimated using CTFFIND4 without accounting for the tilt geometry. Initial ribosome positions and orientations were determined using template matching on 8x binned tomograms, resulting in approximately ∼240K particles. These were then iteratively aligned at 8x and 4x using a mask shaped to contours of the full ribosome density. Particle scores were distributed bimodally; the ∼100K particles in the higher-scoring distribution were selected for further processing. These particles were further aligned at 2x binning, first using a full ribosome mask, followed by alignment using a mask focused on the large subunit. Tilt-corrected periodogram averaging was performed using tiltCTF module in TOMOMAN. 3D-CTF corrected tomograms were reconstructed using novaCTF module of TOMOMAN, using the new CTF parameters. Particles were extracted using the particle list generated with tilt uncorrected tomograms at 2X binning. These particles were further aligned using same steps as before at 2x binning, first using a full ribosome mask, followed by alignment using a mask focused on the large subunit.

